# Linking hubness, embryonic neurogenesis, transcriptomics and diseases in human brain networks

**DOI:** 10.1101/2022.04.01.486541

**Authors:** Ibai Diez, Fernando Garcia-Moreno, Nayara Carral-Sainz, Sebastiano Stramaglia, Alicia Nieto-Reyes, Mauro D’Amato, Jesús Maria Cortes, Paolo Bonifazi

## Abstract

Understanding the architectural principles that shape human brain networks is a major challenge for systems neuroscience. We hypothesize that the centrality of the different brain circuits in the human connectome is a product of their embryogenic age, such that early-born nodes should become stronger hubs than those born later. Using a human brain segmentation based on embryogenic age, we observed that nodes’ structural centrality correlated with their embryogenic age, fully confirming our hypothesis. Distinct trends were found at different resolutions on a functional level. The difference in embryonic age between nodes inversely correlated with the probability of existence of links and their weights. Brain transcriptomic analysis revealed strong associations between embryonic age, structure-function centrality, and the expression of genes related to nervous system development, synapse regulation and human neurological diseases. Our results highlight two key principles regarding the wiring of the human brain, “preferential age attachment” and “the older gets richer”.

## MAIN TEXT

The most characteristic anatomical property of brain networks is their organization across multiple spatial scales. Micro-scale neurons are interconnected within local circuits or microdomains, which are scaled up to the macro-scale over which long-range connections allow distant neurons to communicate and different circuits to interact(^1, 2^).

Consequently, a key challenge in fundamental and clinical neuroscience is to decipher the rules of connectivity that shape brain networks in order to understand how the brain works and how traumatic or neurological damage may affect brain functionality(^3^).

The general structure and function of the human brain, and its internal connectivity are all the result of its developmental history, which is at the same time the product of evolution(^4^).

As such, the brain follows developmental instructions to establish a blueprint of diverse neural connections, the functioning of which can be later refined.

Previous studies have evaluated the development of brain networks at different stages of maturation, particularly in relation to the appearance of key network structures like central nodes(^5^). However, there are still no detailed studies that fully relate the connectivity of the adult human brain with the sequential (evolutionarily preserved) neurogenesis of circuits.

Complex networks have emerged over the past two decades and provide a powerful mathematical framework to quantify the complexity of brain circuitries.

Different studies that assessed the data from real-world networks’ have highlighted that small-world, scale-free or heavy-tailed distribution network organizations are stereotypic topologies of many distinct domains(^6, 7^).

These key topologies have been identified at the structural-functional level in microcircuits, and in meso- and macro-scale networks(^8^).

A pioneering study(^9^) showed that the principle “the rich gets richer” (a.k.a. “preferential attachment”) led to the development of the scale-free networks that are characterized by highly connected rare nodes or hubs.

Inspired by this model(^9^), here we aim to combine the heavy-tailed signatures widely observed in brain networks’ with the stereotypic, evolutionary-preserved sequential neurogenesis in the developing brain. In this way, we will test the hypothesis that the topology of brain networks could be shaped according to the rule that “the older gets richer”, i.e. the evolutionary older circuits or those generated earlier in embryogenesis are those most central in the organization of the adult brain network(^10^).

As a consequence, quantification of the hubness of brain circuits based on metrics of centrality in complex networks should be correlated with embryogenic age. Our hypothesis extrapolates on the macro-scale the earlier pioneering evidence that GABAergic neuronal hubs were functionally identified in hippocampal circuits on a micro-circuit level, based on early born GABAergic neurons in both developing and adult murine circuits(^11–13^).

To test our hypothesis, we differentiated and segmented human brain circuits according to their *First neurogenic Time (FirsT)* following the criteria established in the methods, i.e. the post-conception day (or embryonic day) on which the first neurons of the circuit are generated in the corresponding developmental neuromere (Fig. 1A). The match between developmental neuromeres and adult circuits, and the estimation of FirsT in humans extrapolated from different animal species, is presented with underlying literature and full references in the Method section. As such, a list was generated that organized the human circuits according to their FirsT, from older to younger circuits (Table 1). We identified 18 distinct macro-circuits (MACs) corresponding to 18 separated fundamental developmental units for which a timing sequence based on their FirsT could be defined. Other relevant circuits for which detailed developmental information is known could not be segmented on MRI, such as the neocortical layers and thalamic nuclei (see Supplementary Information).

**Figure 1.**
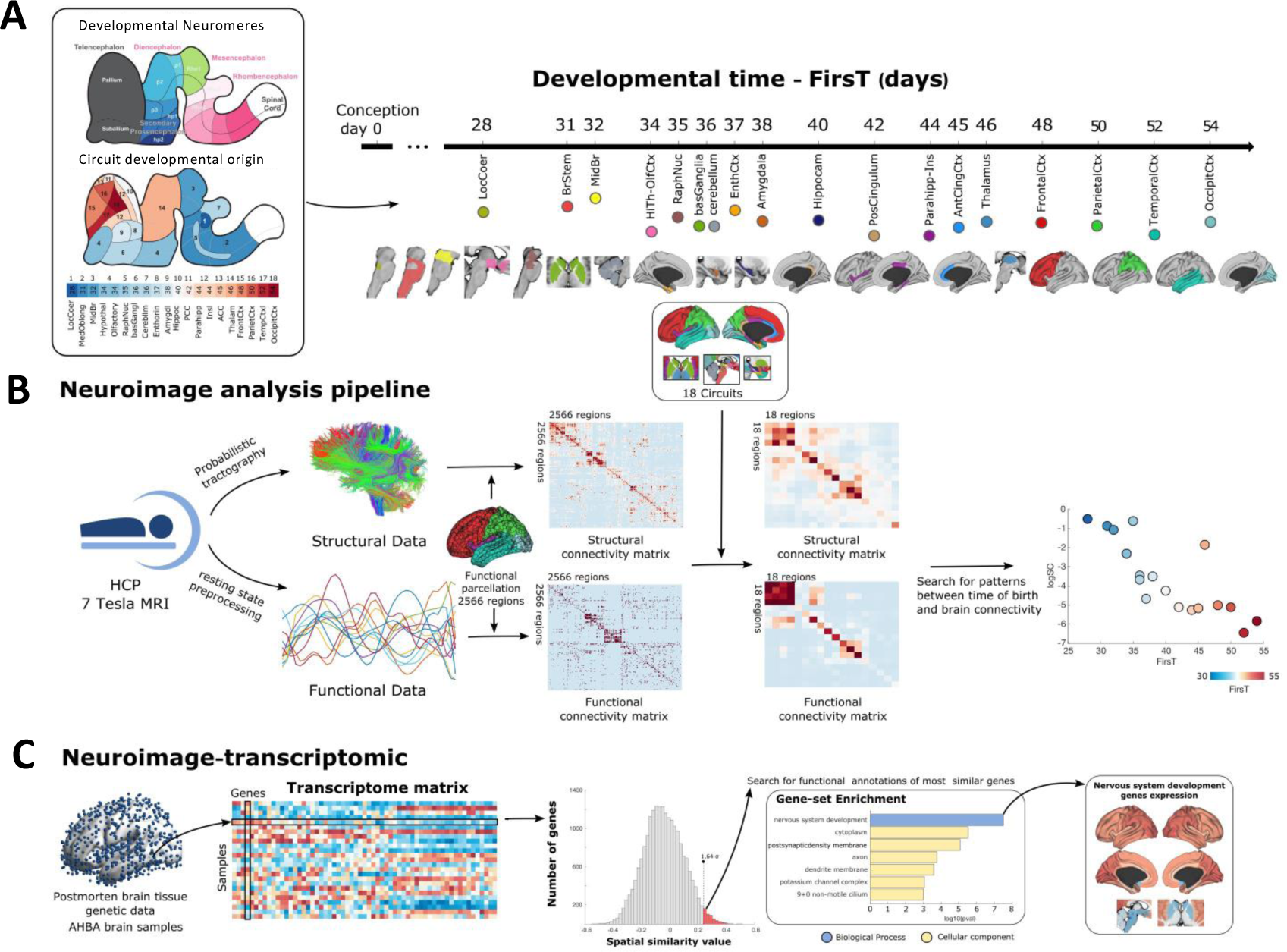
From the circuits’ embryogenic age to brain networks and transcriptomics. (A) Sagittal schemes of the early embryonic human brain. The upper scheme represents the neuromeres and fundamental regions of development, the lower scheme displays the location of the 18 MACs within the early embryonic human brain, colour-coded according to their FirsT. (B) Scheme of the neuroimage analysis pipeline. Brain networks were reconstructed from healthy adult subjects scanned by MRI at 7T within the HCP initiative. For each subject, the structural and functional networks were reconstructed using probabilistic tractography and resting-state activity, respectively. For every pair of ROIs (2,566 ROIs), the structural connectivity matrix represents the putative number of fibres connecting them, while the functional connectivity matrix reports the correlation in their activity as revealed by the BOLD time series. The averages across subjects were used as final representative brain networks (see Methods). From the 2566 ROIs’ matrices, the 18 MACs’ networks were reconstructed and represents for each pair of MACs the average links between the ROIs forming them (see Methods). To identify patterns between networks’ topology and neurogenesis, the correlation between the nodes’ metrics (centrality or segregation) and embryonic age was calculated. (C) Brain transcriptome data from AHBA dataset (see methods) was used to search for protein coding genes with a high similarity between its spatial brain expression and embryonic age. Functional annotations of the obtained genes were further computed using overrepresentation analysis to find significantly associated biological processes and cellular components associated with embryonic age.

**Table 1.** Developmental information of the human brain macrocircuits studied. The table includes the list of 18 MACs, the developmental brain region from which each MAC is derived, the main anatomical brain structures contained in each MAC, information about the earliest stage at which neurogenesis (FirsT) is observed or predicted in the human brain, and in other mammalian species, and the number of ROIs in each MAC. References in Supplementary Material.

|  | Major developmental brain area | Circuit | fMRI Comprised structures | Translated earliest stage in humans (embryonic days) | Earliest stage in other species | source references | Observations | ROIs # |
| --- | --- | --- | --- | --- | --- | --- | --- | --- |
| 1 | Rhombencephalon | Locus Coeruleus | Locus Coeruleus | 28 | Rat - Soon before E12; Mouse - E9 | T1–4 | Likely one of the earliest generated nucleus. Noradrenergic neurons are the earliest-born within the LC | 2 |
| 2 | Rhombencephalon | Medulla Oblongata | Pons and Medulla oblongata | 31 | Rat - Soon after 12 | T1 | Brainstem neurogenesis starts much earlier than in the rest of the brain. However, we could not segment its many different compartments by fMRI and so it appears fused in one single circuit. | 40 |
| 3 | Mesencephalon | Midbrain | Superior colliculus, inferior colliculus, tegmental midbrain | 32 | Mouse - E10.5 | T5,6 | Substantia nigra pars reticulata is the earliest mesencephalic nucleus to start neurogenesis | 23 |
| 4 | Secondary Prosencephalon | Hypothalamus | Hypothalamus, mammillary bodies | 34 | Rat - E12; Macaque - E33; Cat - E21 | T7–9; translatingtime.org | Lateral hypothalamic nuclei (lateral hypothalamic nucleus, lateral preoptic area and lateral mammillary bodies) start hypothalamic neurogenesis at | 18 |
|  | Telencephalon | Olfactory | Olfactory Cortices | 34 | Mouse E11 | T10; translatingtime.org | Unique segmentation and ROI identification with Hypothalamus |  |
| 5 | Rhombencephalon | Raphe Nuclei | Raphe nuclei | 35 | Rat E12 | T1 | Neurogenesis in the Raphe nuclei starts in the Raphe Magnus nucleus | 3 |
| 6 | Telencephalon | Basal Ganglia | Caudate, Putamen, Accumbens, Septum | 36 | Mouse - E11; Macaque - E38 | T11 | Accumbens starts neurogenesis at E38 in macaque; Striatum and globus pallidus start soon after. | 77 |
| 7 | Rhombencephalon | Cerebellum | Cerebellar cortex, deep cerebellar nuclei | 36 | Mouse E11; Rat - E13 | T12–14 | Earliest neurons are projection neurons of the deep cerebellar nuclei, followed by Purkinje neurons. | 275 |
| 8 | Telencephalon | Entorhinal | Entorhinal cortex | 37 | Rat - soon after E14 | T15 | Neurogenesis starts earlier to other parahippocampal cortices | 11 |
| 9 | Telencephalon | Amygdala | Pallial amygdala, subpallial amygdala | 38 | Mouse E11 | T16,17 | Neurogenesis starts with GABAergic populations of central and medial nuclei, followed by the glutamatergic neurons of the baso-lateral complex | 8 |
| 10 | Telencephalon | Hippocampus | Amon horn, Dentate gyrus | 40 | Rat E14.5 | T15 | Earliest neurons occupy the superficial and deep layers of the Ammon's horn (CA1 to CA3), and the hilus | 21 |
| 11 | Telencephalon | Posterior cingulum | Posterior cingulate cortex | 42 | Rat E14.5 | translatingtime.org | Inferred from translatingtime.org Earlier than the anterior cingulate cortex | 22 |
| 12 | Telencephalon | Parahippocampal cortices | Parahippocampal cortex | 44 | Rat E15 | T15 | Includes subiculum and parasubiculum. Unique segmentation and ROI identification with Insula | 107 |
|  | Telencephalon | Insula | Insular cortex | 44 | Rat E15 | T18 | First insular neurons occupy the deep layers of the ventral agranular insular cortex. |  |
| 13 | Telencephalon | Anterior Cingulate cortex | Caudal Anterior Cingulate, Rostral Anterior Cingulate | 45 | Macaque - E40 | T19 | First Layer VI neurons in the anterior cingulate cortex were revealed with [3H]-thymidine injection at E40 in the macaque brain | 37 |
| 14 | Diencephalon | Thalamus | Thalamic nuclei: LP, VL, VPL, LGN, MGN, Pulvinar, | 46 | Rat - E14 | T20 | Several thalamic nuclei (VB, VE, LGN, PO, MGN and LP) start neurogenesis at E14 in the rat brain. | 28 |
| 15 | Telencephalon | Frontal cortex | Caudal middle frontal, lateral orbito frontal, medial orbito frontal, paracentral, pars opercularis, pars orbitalis, pars triangularis, precentral, rostral middle frontal, frontal pole and superior frontal cortices | 48 | Macaque - E40; Ferret - E30; Mouse - E11.5 | T21–23 translatingtime.org | Neurogenesis in the cortex starts synchronously throughout all cortical areas. The earliest-generated cortical populations are the Cajal-Retzius cells of the marginal zone and the subplate neurons of layer 6b. Most of these neurons die early after birth, which is why we considered the early generation of deep cortical layers as the onset of a functional cortical neurogenesis. In these layers, there is a gradient of neurogenesis starting first at the rostral pole of the cortex, and finishing last at the caudal regions. In macaque, frontal cortex starts at E45, and occipital cortex starts at E54. In human, we estimate this gradient as a 2-day difference between cortical lobes. | 719 |
| 16 | Telencephalon | Parietal cortex | Inferior parietal, postcentral, precuneus, superior parietal and supramarginal cortices | 50 |  |  |  | 502 |
| 17 | Telencephalon | Temporal cortex | Banks of the Superior Temporal Sulcus, fusiform, inferior temporal, superior temporal, and transverse temporal cortices | 52 |  |  |  | 384 |
| 18 | Telencephalon | Occipital cortex | Cuneus, Lateral occipital cortex, lingual cortex, pericalcarine cortex | 54 |  |  |  | 289 |

We used dMRI and resting-state fMRI images acquired at 7T from N=184 healthy subjects taken from the Human Connectome project to reconstruct structural-functional brain networks (see Fig. 1B and Methods). Since the volume of the different MACs spanned more than two orders of magnitude (see Supplementary Fig. 1B), the MACs were also parcellated (see Methods) to obtain spatially segregated ROIs with a volume comparable to the smallest circuits (e.g., the Locus Coeruleus -LC), each in the range of several dozen voxels (see Supplementary Fig. 1A for the overall ROI volume distribution and Table 1 for the number of ROIs per MAC).

Each of the 18 MACs had a corresponding FirsT, which was also assigned to all ROIs obtained for a given MAC, and a total of 2,566 ROIs were defined in the brain. To explore the possible existence of patterns of correlation between FirsT and brain circuit connectivity at different spatial resolutions, eigenvector centrality (EC) was calculated for each ROI and MAC (Fig. 1B; see Methods for the rationale behind the choice of metrics). In addition, the brain maps of the transcriptome of 20787 coding-genes were related to the brain maps of neurogenesis (FirsT) and nodes’ hubness (Fig. 1C) in order to obtain a complementary biological correlate to the brain connectivity model.

To quantify the centrality of each ROI and MAC in the brain networks, we calculated EC on high spatial resolution networks composed of the 2,566 ROIs (referred to as high-resolution structural and functional connectivity networks, SC^HR^ and FC^HR^) and low spatial resolution networks composed of the 18 MACs (referred to as low-resolution structural and functional connectivity networks, SC^LR^ and FC^LR^).

The high-resolution networks were calculated from the average of all the subjects, based on probabilistic tractography and resting-state activity correlations (note that only positive functional links with significant correlations, passing the Bonferroni multiple-comparison correction, were considered; see Methods). The low-resolution networks were calculated as the average of the links connecting the ROIs within the MAC pairs in the SC^HR^ and FC^HR^. Note that while the diagonal of the connectivity matrices (self-connections) was taken to be zero in the high-resolution space, in the low-resolution space it was not zero but rather, it represented the average strength (defined as the sum of links’ weights) of the ROIs within a given MAC (internal MAC connectivity, Supplementary figure 3).

To test our main hypothesis, i.e. that the structural centrality of brain nodes is inversely correlated with embryogenic age, such that the earlier the FirsT the higher the centrality, we computed the Spearman’s partial correlation (SP-pCorr) between the EC and FirsT, eliminating the effect of the cofounder white matter volume per ROI or MAC that also shows a gradient between early and late generated structures (Supplementary Fig. 2). Similarly, we repeated the same calculation for the functional networks, in this case removing the cofounder of grey matter volume per ROI or MAC (Supplementary Fig. 2). The plots of the EC obtained for the SC and FC are summarized at both spatial resolutions in Fig. 2A.

**Figure 2.**
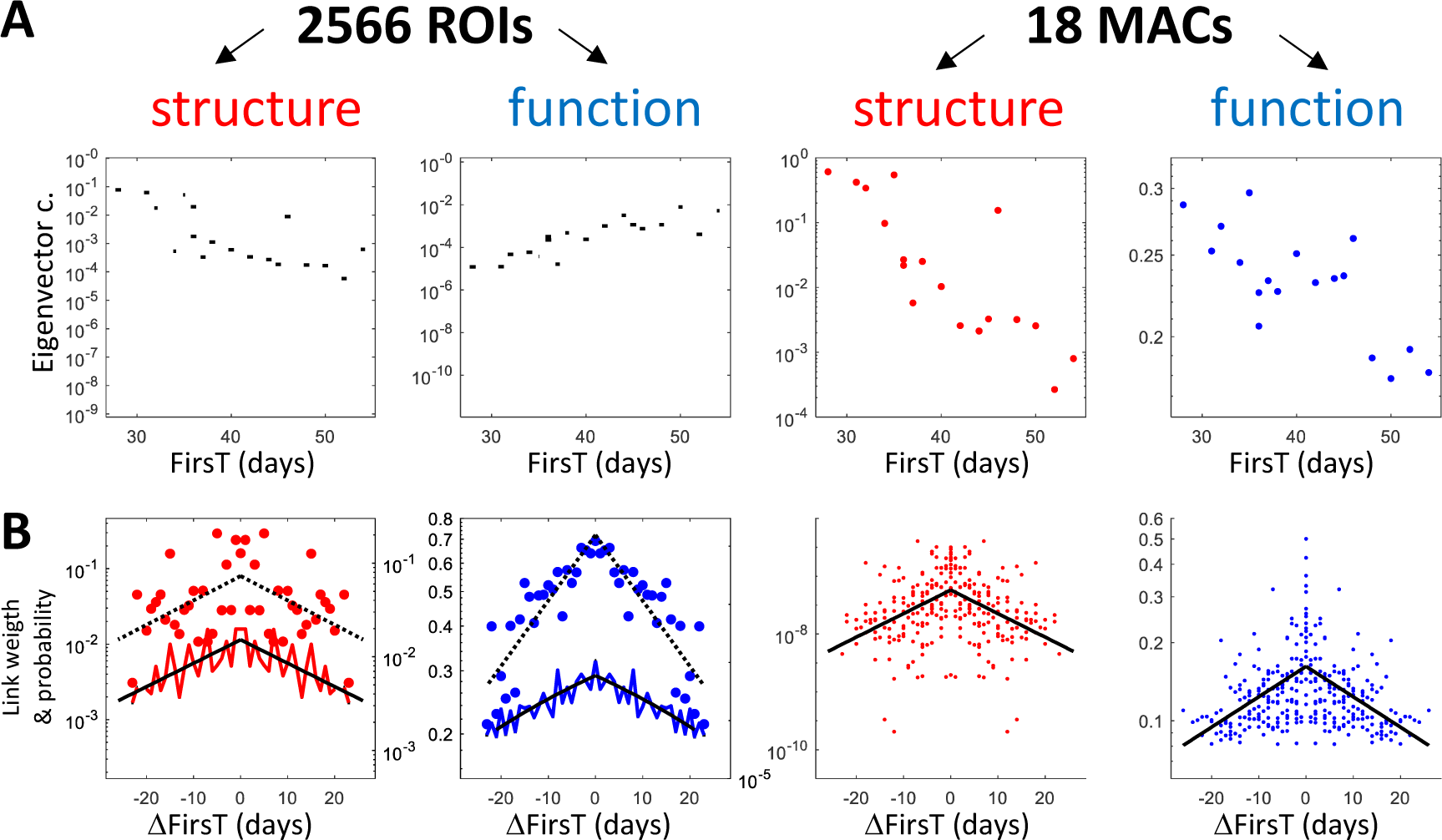
Scatter plots of the nodes’ centrality and the links’ probability/weight relative to the FirsT, in the high and low resolution structural-functional networks. (A) Scatter plots of the nodes’ centrality as a function of embryonic age. Red and blue indicates the structural and functional cases, respectively. The first and second columns from left represent the results for the 2,566 ROIs as violin plots, where the mean of each group (i.e. ROIs within a MAC) is plotted in black, and the third and fourth columns represents the 18 MACs. All plots have a logarithmic scale in the y-axis. (B) Scatter plots of the link weights (left y-axis) as a function of the differences in the first neurogenic birthdate. The colours and columns are as indicated in panel A. The violin plots for the 2,566 ROIs show the link weight distributions while the solid blue/red lines plot the average values. Black solid lines show the exponential (linear) fit on the average values for the structural (functional) case. The link probability is represented by the right y-axes and displayed as dots, and the broken black line showing the exponential fit. All plots have a logarithmic scale in the y-axis.

When looking at the high-resolution networks, the centrality of the SC^HR^ and FC^HR^ nodes showed significant trends with FirsT, respectively with negative (SP-pCorr=-0.44, p=0.001) and positive (SP-pCorr=0.34, p=0.01) correlations. Note that the negative correlations observed for the SC^HR^ (physically constrained spatial domain) support the leading hypothesis that early born brain regions (e.g., the LC and brainstem) have greater “hubness” or centrality, we refer to this as “the older gets richer” principle. By contrast, in the case of the FC^HR^ (functional domain related to time and dynamics) the positive correlations observed between centrality and FirsT highlight the greater functional centrality of late brain circuits (such as neocortical ones).

For low-resolution networks, we observed negative centrality correlations with respect to FirsT in both the SC^LR^ (SP-pCorr =-0.81, p<0.001) and FC^LR^ (SP-pCorr =-0.63, p=0.007). The latter result shows a clear inverted trend between functional networks in low and high-resolution. The inverted trend between the time of birth of the 18 different MACs and the EC of the SC^LR^ and FC^LR^ can be visualized on a brain surface (Fig. 3A). In order to understand the inversion of the EC trend between the FC^LR^ and FC^HR^, we first dissected the contribution of the links between the ROIs belonging to the same MAC (since these are not considered by the centrality measure in the low-resolution case; we refer to them as internal links, *int-Links*, see Methods), by looking at the ROIs’ strength (i.e. the sum of the links’ weights of the ROIs). In the high-resolution networks, the trend of the average ROIs’ strength as function of FirsT (Supplementary fig. 3A) did not change when the *int-Links* were discarded, and it displayed similar trends as in the case of the ROIs’ EC shown in fig. 2A. Focusing next on the MACs’ links (Supplementary fig. 3B) as function of FirsT, we observed the presence of few but very strong links in earlier MACs, which also displayed overall higher strength, with explain the negative trend between MACs’ strength and FirsT as observed for the EC.

**Figure 3.**
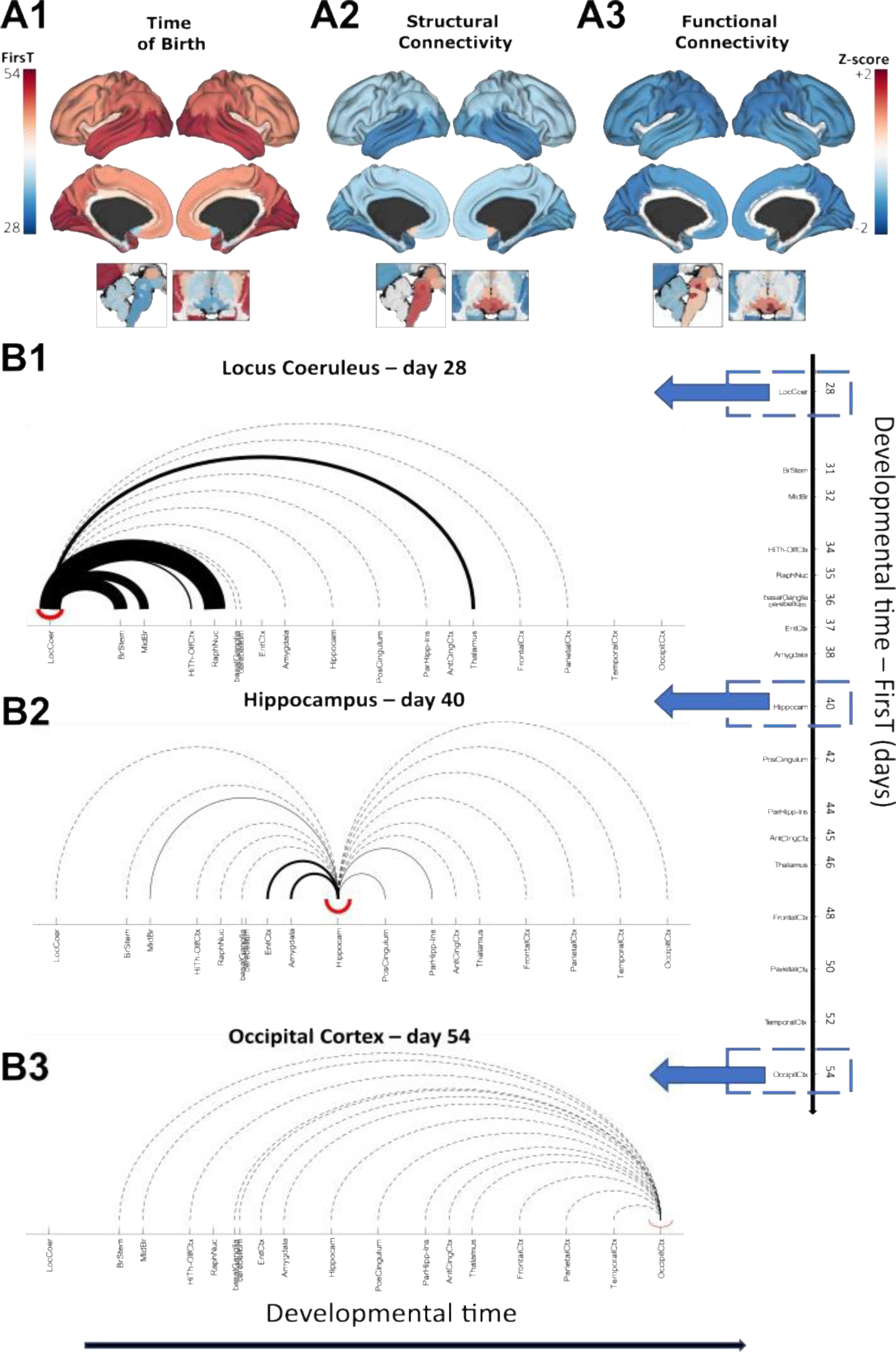
Visualization of brain connectivity and the first neurogenic birthdate. (A) The colour coded glass brain maps of the eighteen MACs represents the FirsT (A1; see colour bar on the left) and the z-score (see colour bar on the right) of the eigenvector centrality for the SC^LR^ (A2) and the FC^LR^ (A3). (B) Linear temporal representation with arches of the MACs structural connectivity for the earliest (B1) generated MAC (Locus Coeruleus, FirsT 28), for a representative mid-term (B2) generated MAC (Hippocampus, FirsT 40) and for the latest (B3) generated MAC (Occipital Cortex, FirsT 54). The x-axis represents the developmental time of several FirsT, where the different MACs are located according to their FirsT. Note that the Basal Ganglia and Cerebellum overlap since their neurogenesis is initiated on day 36. The thickness of top black arches is proportional to the link weight between the MACs extracted from the 18 node adjacency matrix. Broken black lines mark the presence of connections but with little weight (i.e. below the lowest threshold for the visualization of solid lines).

We next checked whether the difference in the time of neurogenesis (ΔFirsT) might also be related to the probability of the existence and the weight of the links (Fig. 2B). In the SC^HR^ and FC^HR^, where a connection was present in 5% and 6.5% of the overall node pairs respectively, the link probability decreased as a function of ΔFirsT with a Spearman correlation of -0.53 (p<0.01) and -0.93 (p<10^-10^). Similarly, the link weight decreased with a correlation of -0.81 (p<0.001) and -0.87 (p<10^-5^). In the SC^LR^ and FC^LR^, only the link weights were considered as these matrices are dense and it decreased as a function of ΔFirsT with a correlation of -0.50 (p<0.001) and -0.52 (p<10^-12^). The plots of the link weights (see Supp. Fig. 4 and Supp. Fig 5 for each MAC in which each plot represents the links of a given MAC and its FirsT is highlighted as a vertical broken line) provide a visualization of how the brain circuits generated at similar neurogenic time points tend to connect more strongly, we refer to this as the “preferential age attachment” principle. As an example, the structural connectivity of three representative MACs was visualized over a linear time scale, relative to the earliest (Locus Coeruleus, 28 days), latest (occipital cortex, FirsT 54 days) and intermediate FirsT (hippocampus, 40 days: Fig. 3B1-3). This visualization shows that the links’ weights of the earliest MAC are much higher than that of the later MACs, and for the case of the LC and hippocampus, it also shows that brain circuits generated at a similar neurogenic time point are more strongly connected.

**Figure 4.**
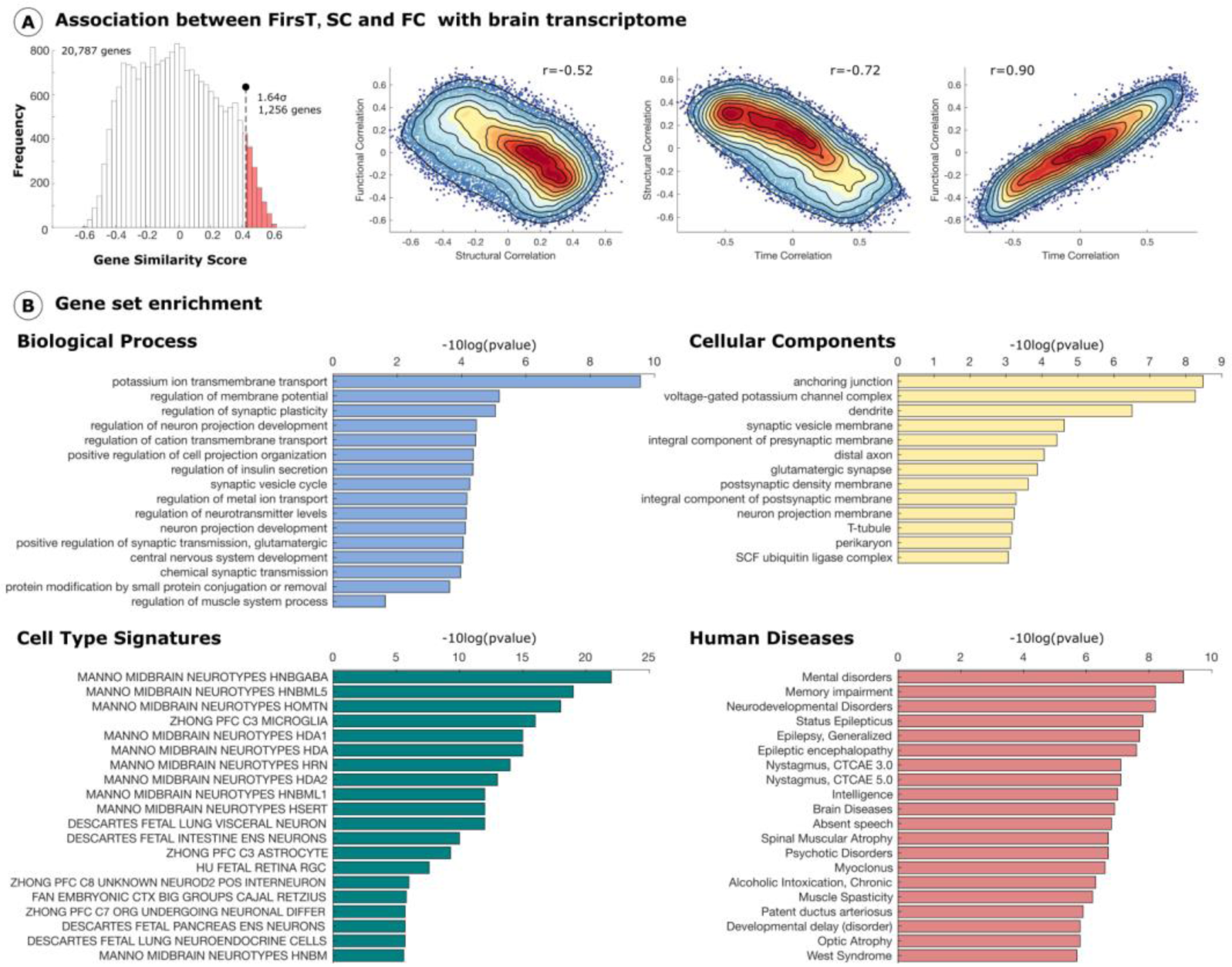
Transcriptome of brain circuits according to embryogenic age. (A) Mean spatial similarity between the expression of 20,737 protein-coding genes with the time of birth and structural and functional centrality. Structural and functional centrality of the 2,566 brain regions was downsampled to 90 region (with gene expression data) and for each of the 90 regions one of the 18 FirsT was assigned. (B) 1,256 genes from the positive tail (6% of protein coding genes) – genes whose spatial expression is most similar to FirsT and functional and structural centrality – were used to compute the overrepresentation analysis for different annotations: biological processes, cell components, cell types, and human diseases.

**Figure 5.**
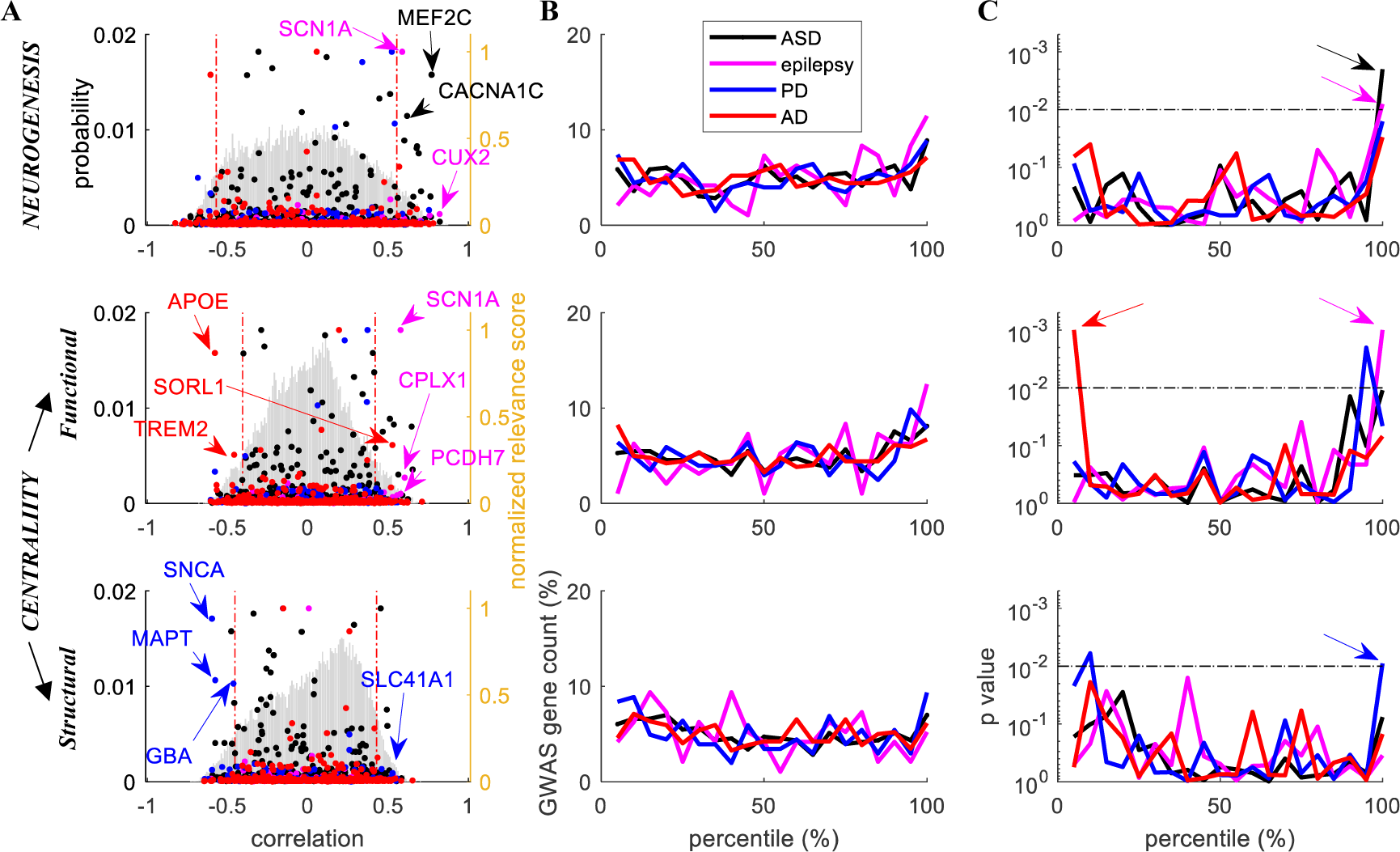
Correlations between the maps of the gene transcriptome, neurogenesis and hubness in relation to Autism Spectrum Disorder, epilepsy, Parkinson’s and Alzheimer’s disease. For each gene the spatial expression in the brain was correlated to the spatial maps of eigenvector centrality and neurogenesis (note that from the sub-sampled brain transcriptomes, 90 brain regions were considered: see Methods). **A** Probability distribution of the correlations (light gray histogram, left axis) between transcriptomics (20,787 genes) and neurogenesis (top plot), functional centrality (middle plot) and structural centrality (bottom plot). The normalized relevance score of the GWAS genes listed for each disease (obtained from the GeneCards databases; Note that normalization is computed separately for each disease; see Methods) has been plotted with dots (right axis). The colors used to display the results for the different diseases in pnale A-C, are shown in the inset legend of the top plot of panel B. Vertical broken lines highlight the lowest and top five percentile interval. Coloured labels and arrows mark the most relevant representative GWAS genes with correlations in the highlighted extreme percentiles.**B** For each disease, the normalized count (percentage of gene per interval) of GWAS genes in five percentile intervals of the correlation distribution (light gray histogram of panel A) is plotted for the different diseases (legend of colours is in the inset of the top plot)). **C** P-value of the normalized gene count in each percentile. The threshold for p<0.01 calculated from the null model (based on a thousand reshuffled replications) which keeps into account spatial dependencies of both centrality maps and gene-expression is plotted as a broken horizontal black line. The arrows mark the extremities of the correlation distributions with a significantly higher number of GWAS-genes than what expected by chance.

To obtain a list of genes potentially associated with the brain circuits’ FirsT and the brain networks’ centrality, post-mortem brain transcriptomic data from the Allen Human Brain Atlas (AHBA) was used. Given the limited spatial sampling across brain regions and donors, we used 90 distinct brain regions with corresponding unique FirsT (supplementary fig. 6), on which gene expression was quantified, and average values of structural and functional centrality was calculated from the englobed ROIs (see Methods). Average Spearman correlations between the brain maps of expression of 20,737 protein coding genes with FirsT, structural and functional centrality was computed (see scatter plots of Fig. 4A). First, the correlation of gene expression with functional centrality displayed a positive linear trend with respect to FirsT (Pearson correlation 0.90, p<0.01), while an opposite trend was observed for structural centrality (Pearson correlation -0.72, p<0.01).This observation shows that the expression of a given gene is similarly correlated to brain maps of FirsT and of functional centrality, in agreement with the fact that FirsT and functional centrality are positively correlated in the high resolution networks. On the contrary, the negative trend observed when gene expression is correlated to FirsT and to structural centrality is also in agreement with the fact that structural centrality and FirsT are inversely correlated. Further confirmation of what was just described is given by the anti-correlation (Pearson correlation -0.52, p<0.01) observed for gene expression in functional and structural centrality maps.

Next, we performed enrichment analysis on the 1,256 genes (6% of the protein coding genes) that were found with a correlation >1.64 standard deviations from the distribution obtained (r>0.43). Gene set enrichment analysis of biological process showed that the genes overrepresented were involved in processes related to development of nervous central system and neuron projection development, and regulation of ion transport, membrane potential, synaptic plasticity and transmission, cell projection organization, insulin secretion, and neurotransmitter levels (Fig. 4B). In addition, different cell components were also overrepresented, such as anchoring junction, voltage-gated potassium channel complexes, dendrites, distal axons, and membranes of synaptic vesicle, pre and post synapses, and neurons (Fig. 4B). Cell type signatures were highly over-represented for several midbrain neurotypes, PFC microglia and astrocytes, and fetal neurons (Fig. 4B). Notably, enrichment of human diseases showed overrepresentation of mental and neurodevelopmental disorders, memory impairment, epilepsy, and other pathologies (Fig. 4B).

Given these final observations in relation to human diseases, next, we tested and dissected the hypothesis that the genetic contribution to neurological diseases is differentially related to the brain networks’ centrality maps and neurogenesis. Therefore, we studied the distribution of the correlation between gene expression, FirsT (fig. 5 top plots), functional (fig. 5 middle plots) and structural (fig. 5 bottom plots) networks’ centrality (we refer to these last two cases as the gex-centrality distributions). We expected that genes causally related to neurological disorders would show extreme correlation values (Fig. 5). In particular, we focused on Autism Spectrum Disorder (ASD), Parkinson’s disease (PD), Alzheimer’s disease (AD) and epilepsy, and we combined two main human gene datasets (see Methods): GeneCards (www.genecards.org) and the Genome-Wide Association Study Catalogue (GWAS Catalogue; https://www.ebi.ac.uk/gwas/). For each gene reported in the GWAS for a given disease, GeneCards dataset provides a relevance score based on the reported literature linking the gene to the disease (coloured dots in Fig. 5A) while statistical evidence of their causal relationship to the risk of developing a disease is guaranteed by the GWAS data. The genes associated with neurological diseases (via GWAS) were not spread evenly across the gex-centrality and neurogenesis distributions (Fig. 5B) and they showed a significant enrichment at the percentile tails (see black arrows in Fig. 5C), supporting our hypothesis. Notably, this observation was typified by genes with a well-known role, such as: *APOE*, *TREM2* and *SORL1* (AD); *SCN1A, CUX2, PCDH7* and *CPLX1* (epilepsy); *SNCA, GBA* and *MAPT* (PD); or *MEF2C* and *CACNA1C* (ASD).

## Discussion

In this work, we use the earliest time of neurogenesis (FirsT) reported for brain circuits to study adult brain networks that were reconstructed from structural-functional (resting-state) MRI with low (MAC level) and high (ROI level) spatial resolution. By correlating centrality metrics of complex networks with the FirsT, we observed a decreasing gradient of structural hubness in MACs and ROIs from those circuits generated earliest to those that appear later. This observation confirms our main working hypothesis that “older nodes get richer” in terms of hubness within brain networks. In the functional domain, the centrality of the MACs follows the same trend, while conversely, the ROIs that developed later have a higher centrality. Since the links present between the ROIs within a MAC do not affect the overall trend of the ROIs strength as function of neurogenesis, we showed how indeed it is the presence of stronger connections between earlier MACs both in the functional and structural case that leads to a negative trend between MACs’ centrality and neurogenesis.

Our results are consistent with brain differences early and late in development. The neurons generated earlier in the human brain mature in an environment favourable to cell growth, migration and axonal pathfinding. By contrast, neurons born later differentiate into a stiffer neuropil that has a dense extracellular matrix and thus, their growth and axonal pathfinding is more severely restricted by prior developmental maturation (^14^). In addition, the growth of the brain itself plays a significant role in the formation of long-range connectivity. Whereas after four of weeks of gestation (GW) when the LC neurons begin their genesis, the brain measures 3-5 mm in length, whereas at eight GW when many cortical neurons are generated, the human foetal brain measures 27-31 mm(^15^). Thus, the formation of general structural connectivity is affected by the timing of neurogenesis.

The “older gets richer” rule stems from the literature on the connectomics of birth-dated populations, both for neuronal populations that we were able to trace through MRI as well as for other smaller regions of the brain that cannot be distinguished at MRI resolution. In the former case, the neurons appearing first in the brain are those of the LC(^16–19^) and the brainstem and midbrain motor neurons, including those of the trigeminal mesencephalic nucleus(^20^). These organize into small nuclei and although most of them cannot be segmented by MRI (here we only considered the LC and Raphe nuclei), our model predicts that they all are potentially relevant in terms of their connectivity. Noradrenergic LC neurons establish connections that span the whole brain(^21^), and both LC and trigeminal nuclei contribute to pathologies such as AD when they degenerate(^22–24^).

Beyond the brainstem, pioneering studies on GABAergic hub cells(^11^) in the developing mouse hippocampus have shown how early born GABAergic neurons operationally and morphologically represent hub cells(^12, 25^), also later identified in other structures(^26^) and in the adult mouse hippocampus(^13^). Within the neocortex, where different neuronal populations are generated over sequential neurogenic stages(^27^) but where these birth-dated circuits cannot be tracked by MRI, hubs are related to neurogenesis. A recent computational study of neocortical circuits reported that targeted injury to layer 5 hub neurons produced the greatest damage to the structural-functional integrity of neocortical circuits(^28^). Significantly, these layer 5 neurons are amongst the earliest neurons to be generated in the neocortex. Neuronal types generated earlier in the hippocampus, cortex, brainstem and mesencephalon might also adopt a hub role due to the timing of their neurogenesis. For example, stimulation of Purkinje cells in the cerebellum (an early born GABAergic neuronal type in a circuit that develops early) has been shown to inhibit spontaneous hippocampal seizures in a mouse model of temporal lobe epilepsy(^29^).

In terms of the topology of brain networks, we observe a decreasing gradient in the probability and strength of connections as a function of the difference in time of neurogenesis, both in structural and functional networks in the high and low degree of spatial resolution. Therefore, nodes generated at similar times showed a higher probability and strength of connection. This latter observation supports the hypothesis that brain networks follow a “preferential age attachment rule”, whereby nodes are more likely to connect if the time of their neurogenesis differs little. From a biological point of view, this might explain why neurons establish their main connections in a short temporal window of plasticity, named the critical period(^30, 31^), before becoming functional and maturing. Therefore, neurons with an equivalent birthdate tend to share these permissive temporal windows, which allows their mutual/reciprocal connection. In this context, a recent study has shown in the hippocampal network that sequential neurogenesis affects connectivity and coactivity of neurons so that same birthdate neurons joined into cell assemblies ^32^.

The rules of “preferential age attachment” and “older gets richer” extrapolate to the context of brain networks, which will therefore adhere to the pioneering model of Barabasi-Albert on the construction of scale-free networks and the genesis of network hubs.

Brain connectivity, like other biological features, is ruled and limited by natural selection. Due to the intimate relationship between development and evolution, it is likely that the rules that relate neurogenic birth to connectivity are conserved across species. Indeed, previous studies on the development of brain networks in *C. elegans* highlight the early evolutionary appearance of neuronal hubs(^33^). Moreover, neurons linked by long-range connections tend to be generated at around the same neurogenic time and early in development(^33^), reinforcing the concept of a temporal windows of plasticity around neuronal birthdates. Ultimately, all connections are shaped evolutionarily to optimize axon length and the speed of connections(^34, 35^), and other vertebrate brains are likely governed by the same rules of development and connectivity.

Indeed, the brain regions shown here to have the most hubness are highly conserved in the vertebrate brain(^36^), suggesting their crucial importance in all vertebrates. These regions include the posterior regions of the brain, such as the brainstem and mesencephalon, and the cerebellum, which are known to be relatively similar across the vertebrate taxa(^37^). In part, the rule of the older gets richer suggests that ancient circuits -such as those in brainstem that are related to autonomic animal functions and directly involved in the animal’s survival-are more influential in the network and more stable across evolution. For example, the highly conserved neurogenic formation of the cerebellum was recently demonstrated in several vertebrate species(^38^) and its remarkable orchestrating hub role has an impact on epileptic brain dynamics(^29^). At the other end of the spectrum, circuits related to associative tasks, more strongly linked to cognition and human-specific behaviour, are not necessarily conserved and they appear later in neurogenesis. In high-resolution networks we show here how their structural hubness (low) is inversely related to their functional one (high).

In relation to brain transcriptomics, we studied how spatial maps of gene expression in adult brains correlate with FirsT and structural-functional centrality maps. We found that genes whose expression is positively correlated with the centrality of structural circuits tend to show stronger expression in the circuits generated first. By contrast, genes whose expression is anti-correlated with the centrality of structural circuits show stronger expression in circuits in which neurogenesis occurs later. The opposite was observed for functional networks and thus, the transcriptomics of structural and functional networks appears to be driven distinctly by genes whose expression is developmentally regulated. Previous studies into the genetic basis of human brain network structure and function presented converging evidence that anatomical connectivity is more strongly shaped by genetics than functional connectivity (see(^39^) for a review). Here, we add that the developmental sequence (specifically the first time point of neurogenesis) is an additional key variable to interpret the genetic influence on the connectome, providing a key feature linking embryogenesis to the topology of adult brain networks. It also sheds light on the longitudinal reconstruction of developing brain networks, a phenomenon that presents key technical challenges(^5^).

Since enrichment analysis in relation to human diseases suggested a role in mental and neurological disorders, next we explored if genes with a known role in neurological diseases are linked closely to network centrality and neurogenesis with the potential to exert an influence on key network hubs. Interestingly, in a recent similar study on the UK Biobank database focused on the genetic architecture (heritability) underlying the white matter connectome (i.e. solely focused on structural connectivity) ^40^, enrichment analyses highlighted the key role of neurodevelopmental processes including neurogenesis, neural differentiation, neural migration, neural projection guidance, and axon development. In addition, structural connectivity displayed significant associations also to polygenic scores for psychiatric, neurological, and behavioral traits.

In our research, we studied the correlation between FirsT, structural and functional network centrality maps, and the expression of genes associated with epilepsy, Autistic Spectrum Disorder (ASD), Parkinson’s disease (PD) and Alzheimer’s disease (AD), models of neurodevelopmental and neurodegenerative diseases. First, we observed that a significant number of genes linked to ASD and epilepsy show high positive correlations to neurogenesis and functional centrality maps in agreement with the evidence that ASD and several forms of epilepsy (such as Dravet syndrome and infant epilepsy) is linked to late brain developmental processes (^41^) and supporting the idea that ASD and infant genetic forms of developmental epilepsy can lead to functional brain maps’ lesion rather than structural ones (^42^). Distinctly, in our model a significant population of genes related to PD express similarly to structural centrality maps, so highly expressed in early generated structures (i.e. structural brain hubs) and with lower expression in later developed structures. This support the idea that PD is a disease linked to targeted structural lesions such as those ones occurring in the dopaminergic neuronal population in the basal midbrain (which is the third in the rank of neurogenesis). In the case of the AD, we identified a significant high number of genes with anti-correlated expression to functional centrality, so potentially poorly expressed in functional hubs (late structures) and viceversa. Therefore, we conclude that dysfunctional properties of the proteins coded by these genes, maybe also associated to alterations in their expression levels, can have potential high impact in brain hubs at developmental, functional and structural levels and can play a key role in the aetiology and associated dysfunctions of brain pathologies. Highlighting some well-known genes with top correlation ranks (see supplementary tables) in relation to the studied brain pathologies, we found a contribution of four genes known to drive PD: *SNCA(*^43, 44^) that encodes the parkinsonism-associated Lewy body protein alpha synuclein, *MAPT(*^45^) encoding the microtubule-associated protein tau, GBA ^46^ encoding the lysosomal enzyme glucocerebrosidase and SLC41A1 ^47^ encoding a magnesium transporter (a candidate for the causative gene in the PARK16). Together our results indicate that *SNCA*, *MAPT* and GBA are expressed more weakly in nodes with higher structural centrality under physiological conditions and viceversa for SLC41A1 (Fig. 5A). We also found associations for the gene encoding apolipoprotein E, *APOE*, which is directly related to AD, cerebral amyloid angiopathy, PD and other neurological diseases(^48^), and for *TREM2*, a receptor expressed in myeloid cells 2 that is related to microglial biology, a high risk of developing AD(^49^) and prion diseases(^50^). In addition, the *SORL1* gene that encodes the sortilin-related receptor, and that has been associated with early and late-onset AD(^51^), was also found to have a strong positive correlation with functional centrality. In the case of ASD, we highlight the gene MEF2C^52^ which regulates cortical inhibitory and excitatory synaptic balance and CACNA1C coding for a calcium channel linked to selective autophagy and altered axon targeting and behavior^53^. For epilepsy, we highlight the presence of the *SCN1A* gene, which encodes the Alpha 1 Subunit of the Sodium Voltage-Activated Channel, occupying position 144 in correlation with the functional centrality of 20,787 genes considered, and with a known causal relationship to Dravet Syndrome, a rare genetic disease causing one of the most devastating forms of childhood generalized epilepsy(^54^).

Notably, many genes known to underlie neurodevelopmental disorders (as the ones for example highlighted in ^55^) lie in the extreme percentiles of the FirsT or centrality distributions (such as CACNA1A, DEPDC5, SCN8A, and more).

As a result of both human disease enrichment and deeper analyses of four neurological conditions, our results provide support for the hypothesis that neurological diseases may be linked to key topological alterations to brain networks in relation to node hubness(^56^). Thus, altered transcription of genes whose expression correlates closely with network centrality could lead to profound neurological lesions and disease. In this context, stroke and other circuit damage to early-generated brain structures (such as those in the brainstem region) are known to have life-threatening consequences relative to later-generated cortical damage.

## Supporting information

functional centrality AD

FirsT AD

structural centrality AD

functional centrality ASD

FirsT ASD

structural centrality ASD

functional centrality Epilepsy

FirsT Epilepsy

structural centrality Epilepsy

all gene correlations

functional centrality PD

FirsT PD

structural centrality PD

## METHODS

The codes to reproduce the presented analyses will be made available to interested researchers upon a motivated request.

### Neurogenesis timeline of brain circuits – Selection of circuits

To select the circuits to study, we needed to reach a compromise between the conflicting resolution of MRI and the developmental structural units. Certainly, there are hundreds of brain structures and circuits that can be segmented by MRI (cf. the list of areas that were identified in the AAL classification (49) and others (50)), although many of these regions do not develop independently (for example, the many neocortical areas). By contrast, there are brain regions that are developmentally relevant but that cannot be precisely segmented by MRI (e.g. thalamic nuclei). Therefore, the two criteria followed to select the list of brain circuits that ultimately studied were developmental independence and MRI segmentability.

#### Developmental independence

Based on how the brain is generated during embryonic development, we chose brain regions that were derived from independent units of the early differentiated brain, called fundamental morphogenic units (FMUs (51)). Early in development, all vertebrate brains divide into different vesicles: secondary prosencephalon, diencephalon, midbrain, and hindbrain. Each of these vesicles can be segmented into several neuromeres, which are developmentally independent rings of the neural tube destined to form the major brain regions. Each neuromere contains several FMUs along its dorsal-ventral axis (figure 1A) and the neurons within a given FMU are formed mostly by neurogenesis. For example, the m1 neuromere in the mesencephalon gives rise to most of the neurons in the superior colliculus, so we consider the superior colliculus to be a developmental unit. When a FMU generates through development several brain regions that we can resolve through MRI (e.g. the occipital neocortex), we have merged these structures into a single MAC, as those regions are not developmentally independent and their FirsTs are therefore the same. The literature available shows the specific date of birth of the different populations in each developmental unit in several mammalian species (see below).

#### MRI segmentability

The human brain evolved following an unprecedented expansion of the neocortex, which develops from a single developmental unit (52). But this expansion came at the expense of shrinking many brain regions, such that some are so small that they cannot be segmented at MRI resolution. For example, the brainstem and pons are generated from the rhombencephalon, and there are up to 11 rhombomeres, each of which hosts a number of FMUs but none of which can be segmented by MRI. Thus, we identified consistently segmentable brain structures from MRIs. Consequently, no circuit was assigned to two different birth dates. Most brain structures were generated during a neurogenic period but the first neurogenic time (FirsT) was considered to be the first day after which neurons began to appear.

### Neurogenesis timeline of brain circuits - Estimation of FirsT

We hypothesize that the strength of adult connectivity in a given region depends on the developmental stage at which it forms. (^57^). The same method to estimate human FirsT was applied for all circuits, this is a summary of the pipeline:

1. Experimental birthdating of the neurons of a given brain area through DNA-synthesis labelling (via BrdU or ^3^H-Thymidine administration) in non-human species (references in Table 1).
2. Determination of the developmental stage at which the first neurons of that brain region were generated in that species (stages on Table 1).
3. Translation of this non-human stage into human developmental stage via translatingtime.org (^57^) which provides the FirsT value of the given brain region. This is possible and reliable thanks to the great conservation of the developmental sequence found across mammals (^57^).

Each FirsT data point is based on at least one article, all articles are referenced in Table 1. When several independent articles displayed research on the same structure (i.e., the locus coeruleus), these led to the same FirsT measurement. Therefore, our FirsT data points represent a single embryonic day as FirsT, not a range/interval.

No single database contains birthdate information for all brain regions examined.

## MRI DATASET AND PROCESSING

### Participants

Neuroimaging data was acquired from the Human Connectome Project (WU-Minn Consortium - PIs David Van Essen and Kamil Ugurbil: 1U54MH091657), funded by the 16 National Institutes of Health (NIH) that support the NIH Blueprint for Neuroscience Research and by the McDonnell Center for Systems Neuroscience (Washington University). For this study, we took data from N = 184 healthy subjects acquired at 7 Tesla (72 males, 112 females; 24 participants between 22-25 years, 84 between 26-30 years, 73 between 31-35 and 2 over 36). High-resolution structural T1 images, functional magnetic resonance images at rest (fMRI) and diffusion tensor images (DTI) were used (for more information on the acquisition parameters see Supplementary Methods and the Human Connectome Project documentation - http://www.humanconnectome.org/).

### Image preprocessing

Minimal pre-processed data was downloaded from the Human Connectome(^58^3). T1 structural images were aligned to the anterior and posterior commissures, skull stripped, corrected for gradient distortion and bias fields, and the non-linear transformation to the MNI152 standard space was computed. Using a one-step resampling approach, resting state functional data was corrected for gradient-non-linearity-induced distortion and movement within runs using a rigid body transformation (six parameter linear transformation) and EPI distortion. Both the transformation from the reference image to the structural T1 image, and from T1 to the standard space were also performed through a one-step resampling approach. A subject specific functional data projection to MNI152 was obtained. Diffusion images were normalized for b0 intensity and EPI distortion, movement and eddy currents were corrected, as were gradient non-linearities. The B0 image was co-registered to the subject’s anatomical T1 images. For more details on the minimal pre-processing of the human connectome refer to the original paper(^58^3). To further reduce the noise in the functional data, a general linear model was used to remove linear and quadratic trends, as well as the contribution of movement, the cerebrospinal fluid and white matter signals. Additionally, band-pass filtering (0.01–0.08 Hz) and spatial smoothing with an isotropic Gaussian kernel of 6 mm FWHM was applied.

### Brain macrocircuits (MACs) at low resolution

Several regions from different brain MRI atlases were combined to generate 18 different macrocircuits. The atlases used included the Automated Anatomical Labeling (AAL(49)), Freesurfers Desikan-killiany atlas(54), CIT168 subcortical atlas(55) and locus coeruleus atlas(56). The LC and raphe nuclei were chosen as the functional ROIs that englobed them (for further details see Table 1).

### Brain regions of interest (ROIs) at high resolution

High resolution parcellation was performed after clustering the functional data to study network properties with about 2,500 ROIs. In particular (and similar to(57)), clustering was performed based on temporal correlations between pairwise voxel time series that impose a constraint to ensure spatially contiguous ROIs. This was performed in two stages: 1. clustering at the single subject level; and 2. a second clustering applied to individual subject data. To avoid ROIs with voxels belonging to several circuits rather than performing clustering on the whole brain, this strategy was applied separately for each circuit. Small circuits were neglected and those voxels were inserted into the nearest circuit. After parcellation, we took all ROIs overlapping some of the existing circuits, resulting in 2,566 regions covering the whole brain.

## STRUCTURAL AND FUNCTIONAL BRAIN NETWORKS

### High-resolution

After image processing, the connectivity matrices were obtained at high-resolution, representing two modalities composed of 2,566 ROIs: structural and functional networks. Individual subject functional connectivity matrices were obtained by averaging all the time series of the voxels belonging to each ROI generated, and by calculating the Pearson correlation value between the time series corresponding to each pair of ROIs. The average of the functional matrices across all subjects was then calculated. Only positive links with an average minimal correlation of 0.19 were considered. This value corresponds to a p-value of 0.7 × 10^-9^ when calculated on time series composed by 900 points. Such p-value threshold was chosen in order to apply a strict Bonferroni correction on a 0.01 p-value of multiple comparisons, i.e. divided by the number of links in the adjacency matrix (which correspond to half of the elements in matrix excluding the diagonal considering that adjacency matrices are symmetric by definition in our methodology, i.e. (2566*2566)-2566/2). This final matrix defined the average high-resolution functional connectivity matrix: FC^HR^, and it had a connectivity density of 6.5%.

FSL functions were used to quantify the individual subject structural connectivity matrices. First FSL BEDPOSTX was used to model crossing fibres and subsequently, probabilistic tractography was performed with PROBTRACKX to generate a #ROI × #ROI matrix per subject. Matrices were normalized between 0 and 1 dividing each element by their maximum. The average across subject matrices were finally obtained and only the top 5% of higher weights were considered to create the high-resolution structural connectivity matrix, SC^HR^, consistent with previous studies (58). ROIs’ hubness quantified with eigenvector centrality in the SC^HR^ in the dense (when all links were considered) or sparse (when only top 5% links were considered) adjacency matrix, correlated with a 0.99 value (p<10^-10^) so they did not show significant differences. For both the functional and structural matrices, the principal diagonal elements were set to zero to avoid ROI self-connectivity interactions.

### Low-resolution

Each individual subject functional connectivity matrix, i.e. an adjacency matrix of 2566×2566 elements, was threshold at 0.19 in order to keep only positive significant links as described for the calculation of the average high-resolution functional connectivity matrix FC^HR^. We call the individual high resolution functional matrix as subFC^HR^. Next for each subject we calculated the individual low-resolution functional connectivity matrix (subFC^LR^) as described in the following. Given a pair of macro-circuits, *MAC_i_* and *MAC_j_ (*with *i≠j)*, and the two sets of ROIs belonging to them *R_i_={ROI_i1_, …, ROI_iN_}* and *R_j_={ROI_j1_, …, ROI_iM_}*, where *N* and *M* are the respective number of ROIs, the average of all functional links in subFC^HR^ connecting the two sets defined the functional link between *MAC_i_* and *MAC_j_*, i.e. the low-resolution matrix element 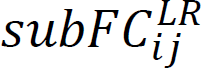 according to:

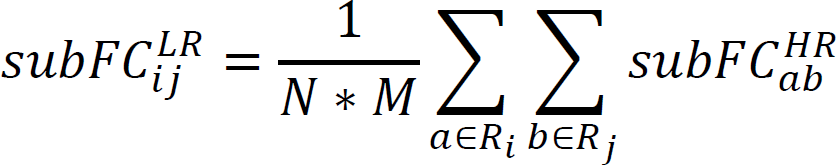

Note that for FC^LR^ the principal diagonal has non-zero elements, representing the internal connectivity within a given MAC, i.e.:

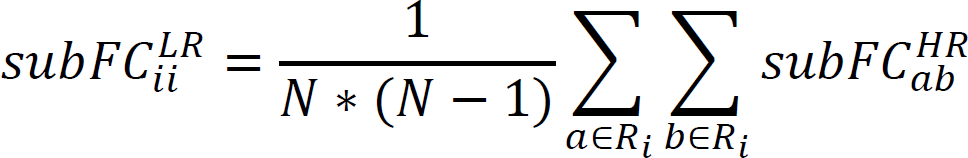

In relation to supplementary figure 3, given a MAC, we call internal links (*int-links*) the subset of links connecting the ROIs belonging to the MAC. Similarly, we define external links (*ext-links*) all the links connecting the same ROIs to the rest of the ROIs in the brain.

In relation to the structural connectivity, we considered the average adjacency high-resolution matrix SC^HR^, and we built the low-resolution structural connectivity matrix, SC^LR^, where the MACs’ volumes (*V_i_* and *V_j_*) were now used for normalization, i.e.:

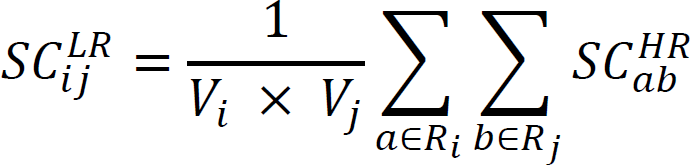

and for the diagonal elements:

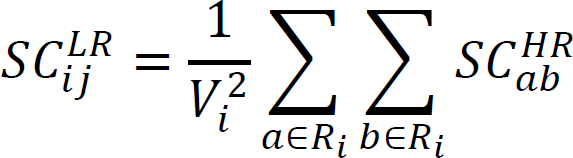

To calculate the *MAC_i_* volume, *V*, we summed the volumes of the corresponding ROIs (set *R_i_*) expressed as the number of voxels.

## COMPLEX NETWORKS ANALYSIS

Through their construction, all the networks analysed were symmetric and weighted, with non-negative values. We used MATLAB (Mathworks) to run different metrics implemented in the Brain Connectivity Toolbox (https://sites.google.com/site/bctnet/measures/list). In particular, we initially chose Node Strength (NS), Eigenvector Centrality (EC), Subgraph Centrality (SC), Betweenness Centrality (BC) and Clustering Coefficient (CC) and we discarded the node degree (as this is not meaningful in fully connected conditions), the flow coefficient (which is not applicable in fully connected conditions), the local efficiency (diverging computational time high number of nodes) and Pagerank Centrality (as this is a close variant of the EC).

Finally, being centrality measures very highly correlated in most networks including the brain^59^ we chose eigenvector centrality as main metric in our study being the one showing less variability when considering all link weights or when applying thresholds to adjacency matrices (as described previously in the Methods at the “high resolution” section). To remove the potential bias introduced by ROIs with a stronger influence of white or grey matter, each individual partial volume of white and grey matter was estimated using the FSL FAST tool(59). For each functional ROI the average white and grey matter partial volume estimates were computed from the group average. We used a Spearman partial correlation to calculate the correlation between structural (functional) centrality and FirsT, while regressing out white (grey) matter partial volume estimates per ROI or per MAC (considering the average across the ROIs within each MAC) in the case of low- and high-resolution networks respectively.

The difference in neurogenesis time between two ROIs or MACs was defined as ΔFirsT. With regards the average connection probability and weight between nodes, as a function of the difference in neurogenesis time ΔFirsT (see fig. 2), the matrix representing ΔFirsT for pairs of ROIs (ΔFirsT^HR^) was first computed according to:

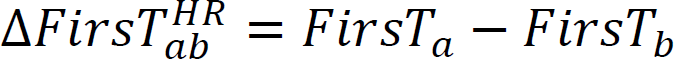

where *(a,b)* is a pair of ROIs.

Similarly, the matrix was calculated for the low-resolution network case for pairs of MACs (ΔFirsT^LR^). Note that by construction, ΔFirsT^HR^ and ΔFirsT^LR^ are antisymmetric. In the case of the high-resolution networks, the Spearman correlation of connection probability and average link weight as a function of ΔFirsT was only calculated considering non-negative elements of ΔFirsT^HR^ and the corresponding elements in the SC^HR^ (FC^HR^: note that the same results would be obtained for the non-positive case), When *F* is the set of elements in the matrix satisfying the condition ΔFirsT^HR^=D (with 0<=D<=max(ΔFirsT)=26 days), the structural (functional) connection probability for ΔFirsT=D was calculated as the fraction of existing links out of the set F in the adjacency matrix SC^HR^ (FC^HR^). Similarly, for the same ΔFirsT=D and set F, the average link weight was calculated from the adjacency matrix SC^HR^ (FC^HR^) for the existing links. In the low-resolution case, since SC^LR^ and FC^LR^ are not a sparse matrix and they have a connectivity density above 95%, only the link weight was calculated as a function of FirsT. As for the high-resolution networks, the Spearman correlation between the link weights and FirsT was calculated on the non-negative links of the SC^LR^ (FC^LR^) and the corresponding elements in (ΔFirsT^LR^).

## GENETIC FINGERPRINT OF BIRTHDATE CIRCUITS

To investigate the genetic fingerprints of the circuits’ birthdays, we used the transcriptome dataset from the AHBA(60,61). The AHBA provides whole-brain genome-wide expression values for 20,737 protein encoding genes extracted from 3,702 brain samples and distributed spatially over six human post-mortem brains. Using the brain sample information, brain maps representing the spatial distribution of each gene in the 18 circuits was generated based on recent recommendations(^60,61^63): i) expression values from multiple probes were averaged; ii) each sample was mapped to one of the 18 circuit atlases, with samples falling outside mapped to the nearest circuit if this was closer than 3 mm; iii) for each individual the median expression values were calculated across all samples that mapped to the same circuit; iv) for each donor a z-score was applied to normalize each gene spatial distribution and minimize inter-subject variation; v) a group expression map was computed by calculating the mean expression values of the six individual donors. While AHBA dataset has not enough spatial coverage for the computation of genetic expression values at each of the 2,566 regions, we generated a 90 regions parcellation to gain more detailed expression maps compared to the 18 regions. The expression atlas containing 90 brain regions was based on the 68 cortical regions of the Desikan-Killiany atlas, the 16 subcortical regions from freesurfer, the cerebellum, brainstem, LC and the dorsal raphe nucleus (see Supplementary figure 6). This 90 regions’ atlas was used to assess the genetic associations of the structural and functional organization of the brain. The centrality of the 2,566 values falling into each 90 regions was averaged to transform 2,566 functional and structural centrality maps to 90 regions.

To search the underlying genetic fingerprints of both the microcircuit FirsT, and the brain functional and structural centrality, a combined score for each of the 20,737 protein-coding genes was computed.

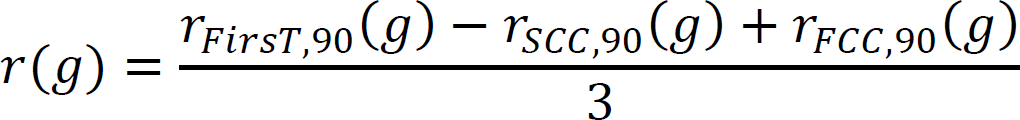

The score *r* for a given gene *g* is the average Spearman correlation between the expression of gene *g* with the FirsT projected to 90 regions *r*_*FirstT*,90_(*g*), with structural connectivity centrality in 90 regions *r*_*SCC*,90_(*g*) and with functional connectivity centrality in 90 regions *r*_*FCC*,90_(*g*). The sign of the Spearman correlation of structural connectivity was inverted as it follows an inverse pattern relative to the FirsT and functional connectivity. Genes with a score >1.64 standard deviations from the distribution obtained were used for further analyses (r>0.43; 1,256 genes).

An overrepresentation analysis was used to find common functional annotations in the obtained gene list. Gene set enrichment was computed using Metascape^62^ (http://www.metascape.org/) for biological processes and cell component annotations from Gene Ontology, human diseases annotations from DisGeNET^63^, and cell type signatures ^62^. Only terms surviving to multiple comparisons are reported; FDR q<0.05. We used Monte Carlo simulation to assess significance of each of the obtained significant annotations, to overcome false-positives bias due to gene co-expression spatial autocorrelation^64^. First, we used BrainSMASH^65^ to generate 1,000 surrogate maps with matching spatial autocorrelation with 90 regions FirsT, and structural and functional centrality maps. Then, for each surrogate map we computed the spatial similarity (Spearman correlation) with 20,737 protein coding genes and we used the list of genes with a similarity >1.64 standard deviations from the obtained distribution to perform an over-representation analysis. For each of the annotations we quantified how many times the enrichment value was higher in the original data compared to the surrogate maps and we generated a p-value of the significance. All the annotations that were not significant (FDR-corrected) were removed.

Regarding genetic and neurological diseases, our analysis was based on the full gene lists related to Epilepsy, ASD, PD and AD as reported by the Genome-Wide Association Study (GWAS) and downloaded from https://www.ebi.ac.uk/gwas/ (GWAS). For these lists of genes, we used the corresponding relevance score for each disease from Genecards database (www.genecards.org). Given a pathology, we normalized the relevance scores dividing by its maximum, in this way we could use a common index wchi could be plot across pathologies (see normalized relevance score in fig. 5) For a detailed definition of the relevance score relative to the disease, see https://www.genecards.org/Guide/Search. The distribution of the genes from the GWAS Catalogue (figure 5) was analysed in intervals of five percentiles. The enrichment for GWAS Catalogue genes in the outermost percentiles vs the rest of the distribution was tested using the previously generated thousand randomizations that preserved both spatial dependencies in the variable representing neurogenesis, functional or structural centrality. In regard to gene expression dependency, the correlation in the expression of the genes in the brain parcellation was preserved in each random trial by reshuffling in the same order the values of the expression across the brain parcellation. Given one thousand randomizations, p-values reported in figure 5 C were calculated with a 0.001 resolution.

The lists of genes from the GWAS list in the outermost percentiles (see vertical black broken lines in Fig. 5A) are reported as supplementary tables, with information on the structural and functional correlations (value, rank and percentile), and the relevance score (value, rank and percentile).

## ACKNOWLEDGEMENTS

We thank M. De Pittá, D. Papo, D. Marinazzo, A. Mazzoni, Y. Ben-Ari for helpful suggestions and critical comments.

## Author contributions

Conceptualization: FGM, PB; Methodology: ID, FGM, NCS, SS, ANR, MDA, JMC, PB; Writing – original draft: FGM, ID, MDA, JMC, PB; Funding acquisition: FGM, JMC, PB.

## ADDITIONAL INFORMATION

Competing interests

The authors declare no competing interests.

## Supplementary materials

Supplementary Information is available for this paper.

Fig S1 – S6

References (T01 to T18)

Auxiliary supplementary tables 01-13

Correspondence and requests for materials should be addressed to Dr. Paolo Bonifazi. Reprints and permissions information is available at www.nature.com/reprints.

