## supplementary figures and supplementary legends for "Linking hubness, embryonic neurogenesis, transcriptomics and diseases in human brain networks"

### EXTENDED DATA FIGURES

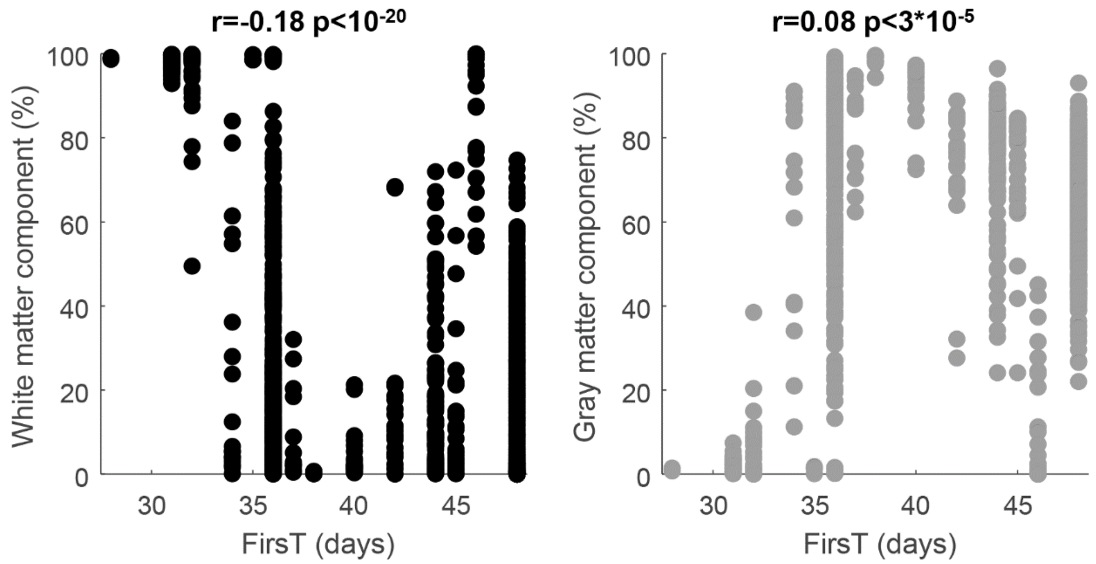


***Supplementary Figure 1. Distributions of MACs and ROIs' volumes. A** distribution of ROIs volume. **B** Cumulative distribution of ROIs and MACs volumes.*

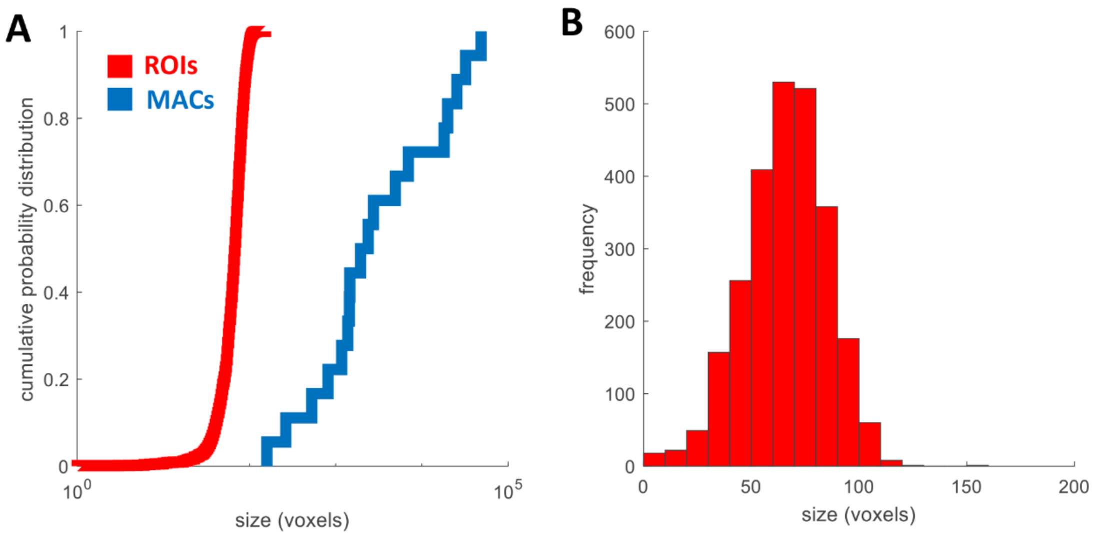


***Supplementary Figure 2.** Percent of white (left plot) and gray (right plot) matter per ROI as function of neurogenesis time.*

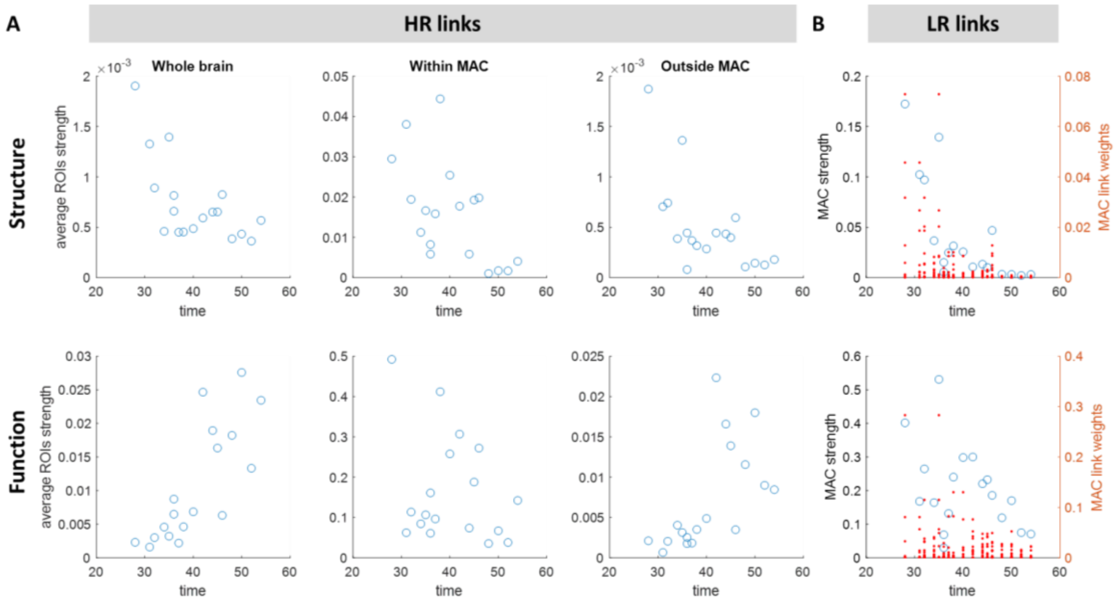


***Supplementary Figure 3. High- and low-resolution nodes' strength: from ROIs' int-links and ext-links to MACs' links and strength. (A)*** *Average ROI strength as function of FirsT. The average is calculated across ROIs with the same FirsT and belonging to the same MAC. The strength is the sum of the links' weight of a given ROI. The links considered are the ones connecting the ROI to: all other ROIs in the brain (int-links plus ext-links; left panels), ROIs belonging to the same MAC (int-links; central panels) and ROIs belonging to different MACs (ext-links; right panels).* ***(B)*** *MACs' strength (blue circles, left y-axis) and weights of MACs' links (red dots, right y-axis) as function of FirsT.*

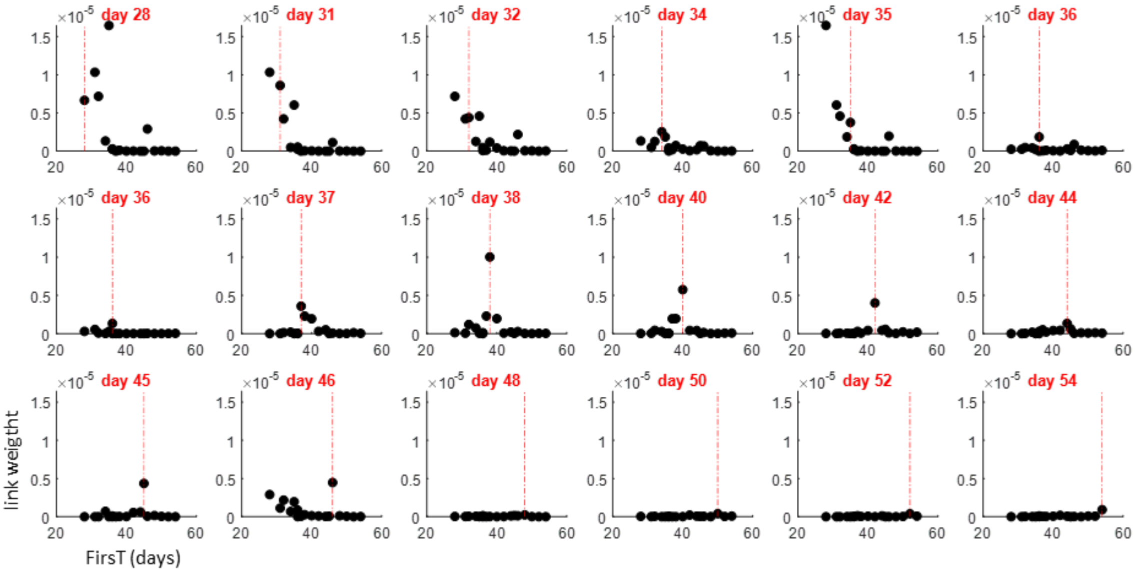


***Supplementary Figure 4.*** *Structural links' weights for each of the MAC according to their time of birth.*

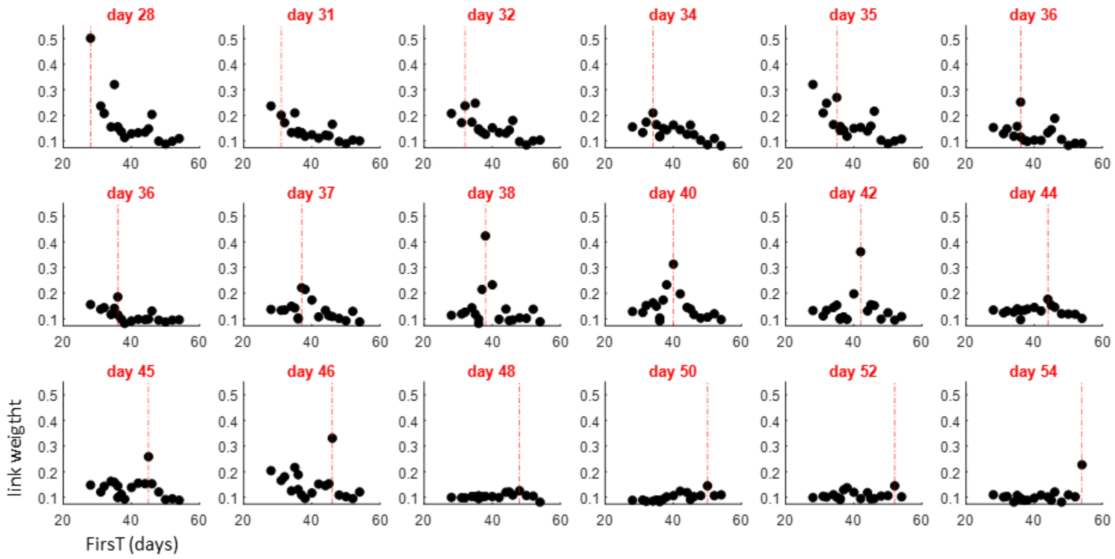


***Supplementary Figure 5.*** *Functional links' weights for each of the MAC according to their time of birth*

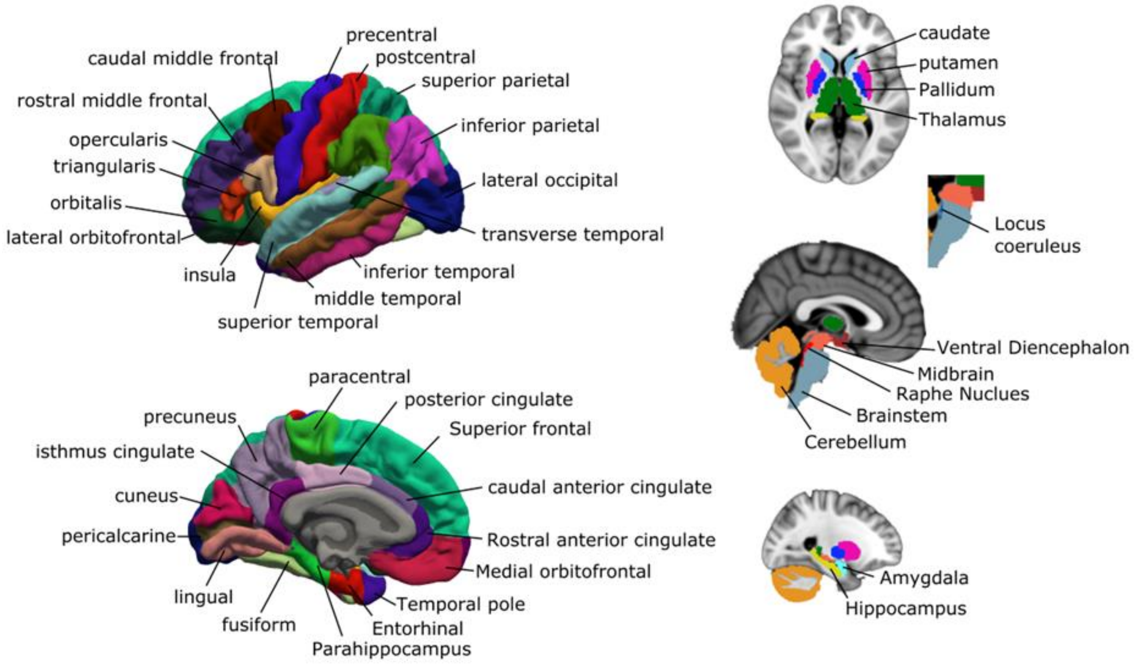


***Supplementary Figure 6.*** 90 Regions parcellation used to project Allen transcriptome database to search for the underlying molecular mechanism behind. The 90 regions atlas includes the 68 cortical regions from Desikan-Killiany atlas, 16 subcortical regions from freesurfer, the cerebellum, brainstem, locus coeruleus and the dorsal raphe nucleus.

***Auxiliary Supplementary tables.***

*The file "gene_correlations_rank_percentiles.xlsx" is the table reporting the 20787 coding-genes with their correlation, rank and percentile in respect to structural and functional centrality, and to FirsT.*

*The other .xlsx files are the tables referring to GWAS genes reported for the four brain disorders considered in this research. Acronyms for diseases and file name are: EPI – epilepsy, ASD – autism spectrum disorder, PD, Parkinson's disease, AD - Alzheimer's disease, FC – functional centrality, SC – structural centrality, FirsT. The tables report the list of GWAS genes related to a given disease, with highest or lowest correlations calculated for FirsT, structural and functional centrality, specifically in the five extreme percentiles of the distributions(see fig. 5A distributions and vertical broken lines). Each table is divided in two parts, the list of genes with highest positive (left side, horizontally highlighted in green on the top) and negative (right side, horizontally highlighted in cyan on the top ) correlations. In the correlation columns (horizontally highlighted in yellow on the top), correlation values, rank of correlation (out of 20787 genes) and percentile of correlation are reported. For simplicity, rank and percentile of positively and negatively correlated genes are calculated from the corresponding extremity of the distribution. In the relevance score columns (horizontally highlighted in orange on the top) Genecards data is considered when available, and accordingly relevance score values, rank (out of the total number of genes in the Genecards list) and percentiles (out of the relevance score distribution for the genes in the Genecards list) are reported.*

### Source references of table 1

*T1. Altman, J. & Bayer, S. A. Development of the brain stem in the rat. I. Thymidine-radiographic study of the time of origin of neurons of the lower medulla. J. Comp. Neurol.* ***194****, 1–35 (1980).*

*T2. Altman, J. & Bayer, S. A. Development of the brain stem in the rat. III. Thymidine-radiographic study of the time of origin of neurons of the vestibular and auditory nuclei of the upper medulla. J. Comp. Neurol.* ***194****, 877–904 (1980).*

*T3. Altman, J. & Bayer, S. A. Development of the brain stem in the rat. V. Thymidine-radiographic study of the time of origin of neurons in the midbrain tegmentum. J. Comp. Neurol.* ***198****, 677–716 (1981).*

*T4. Steindler, D. A. & Trosko, B. K. Two types of locus coeruleus neurons born on different embryonic days in the mouse. Anat Embryol* ***179****, 423–434 (1989).*

*T5. Achim, K. et al. Distinct developmental origins and regulatory mechanisms for GABAergic neurons associated with dopaminergic nuclei in the ventral mesodiencephalic region. Development* ***139****, 2360–2370 (2012).*

*T6. Bayer, ShirleyA., Wills, KatherineV., Triarhou, LazarosC. & Ghetti, B. Time of neuron origin and gradients of neurogenesis in midbrain dopaminergic neurons in the mouse. Exp Brain Res* ***105****, (1995).*

*T7. Altman, J. & Bayer, S. A. Development of the diencephalon in the rat. I. Autoradiographic study of the time of origin and settling patterns of neurons of the hypothalamus. J. Comp. Neurol.* ***182****, 945–971 (1978).*

*T8. Altman, J. & Bayer, S. A. Development of the diencephalon in the rat. II. Correlation of the embryonic development of the hypothalamus with the time of origin of its neurons. J. Comp. Neurol.* ***182****, 973–993 (1978).*

*T9. van Eerdenburg, F. J. C. M. & Rakic, P. Early neurogenesis in the anterior hypothalamus of the rhesus monkey. Developmental Brain Research* ***79****, 290–296 (1994).*

*T10. Workman, A. D., Charvet, C. J., Clancy, B., Darlington, R. B. & Finlay, B. L. Modeling Transformations of Neurodevelopmental Sequences across Mammalian Species. Journal of Neuroscience* ***33****, 7368–7383 (2013).*

*T11. Brand, S. & Rakic, P. Neurogenesis of the nucleus accumbens septi and neighboring septal nuclei in the rhesus monkey: A combined [3H]thymidine and electron microscopic study. Neuroscience* ***5****, 2125–2138 (1980).*

*T12. Miale, I. L. & Sidman, R. L. An autoradiographic analysis of histogenesis in the mouse cerebellum. Experimental Neurology* ***4****, 277–296 (1961).*

*T13. Altman, J. & Bayer, S. A. Development of the diencephalon in the rat. III. Ontogeny of the specialized ventricular linings of the hypothalamic third ventricle. J. Comp. Neurol.* ***182****, 995–1015 (1978).*

*T14. Leto, K., Carletti, B., Williams, I. M., Magrassi, L. & Rossi, F. Different Types of Cerebellar GABAergic Interneurons Originate from a Common Pool of Multipotent Progenitor Cells. Journal of Neuroscience* ***26****, 11682–11694 (2006).*

*T15. Bayer, S. A. Development of the hippocampal region in the rat I. Neurogenesis examined with3H-thymidine autoradiography. J. Comp. Neurol.* ***190****, 87–114 (1980).*

*T16. McConnell, J. & Angevine, J. B. Time of neuron origin in the amygdaloid complex of the mouse. Brain Research **272**, 150–156 (1983).*

*T17. Soma, M. et al. Development of the mouse amygdala as revealed by enhanced green fluorescent protein gene transfer by means of in utero electroporation. J. Comp. Neurol. **513**, 113–128 (2009).*

*T18. Bayer, S. A. Neurogenesis in the rat primary olfactory cortex. International Journal of Developmental Neuroscience **4**, 251–271 (1986).*

*T19. Granger, B., Tekaia, F., Le Sourd, A. M., Rakic, P. & Bourgeois, J.-P. Tempo of neurogenesis and synaptogenesis in the primate cingulate mesocortex: Comparison with the neocortex. J. Comp. Neurol. **360**, 363–376 (1995).*

*T20. Altman, J. & Bayer, S. A. Development of the diencephalon in the rat. IV. Quantitative study of the time of origin of neurons and the internuclear chronological gradients in the thalamus. J. Comp. Neurol. **188**, 455–471 (1979).*

*T21. Rakic, P. Pre- and post-developmental neurogenesis in primates. Clinical Neuroscience Research **2**, 29–39 (2002).*

*T22. Reillo, I. & Borrell, V. Germinal Zones in the Developing Cerebral Cortex of Ferret: Ontogeny, Cell Cycle Kinetics, and Diversity of Progenitors. Cerebral Cortex **22**, 2039–2054 (2012).*

*T23. Charvet, C. J. Distinct developmental growth patterns account for the disproportionate expansion of the rostral and caudal isocortex in evolution. Front. Hum. Neurosci. **8**, (2014).*
